# The Role of Colony Temperature in the Entrainment of Circadian Rhythms of Honey Bee Foragers

**DOI:** 10.1101/2020.08.17.254722

**Authors:** Manuel A. Giannoni-Guzmán, Emmanuel Rivera, Janpierre Aleman-Rios, Alexander M. Melendez Moreno, Melina Perez Ramos, Eddie Pérez-Claudio, Darimar Loubriel, Darrell Moore, Tugrul Giray, Jose L. Agosto-Rivera

## Abstract

Honey bees utilize their circadian rhythms to accurately predict the time of day. This ability allows foragers to remember the specific timing of food availability and its location for several days. Previous studies have provided strong evidence toward light/dark cycles being the primary Zeitgeber for honey bees. Work in our laboratory described large individual variation in the endogenous period length of honey bee foragers from the same colony and differences in the endogenous rhythms under different constant temperatures. In this study, we further this work by examining temperature inside the honey bee colony. By placing temperature and light data loggers at different locations inside the colony we measured temperature at various locations within the colony. We observed significant oscillations of temperature inside the hive, that show seasonal patterns. We then simulated the observed temperature oscillations in the laboratory and found that using the temperature cycle as a Zeitgeber, foragers present large individual differences in the phase of locomotor rhythms with respect to temperature. Moreover, foragers successfully synchronize their locomotor rhythms to these simulated temperature cycles. Advancing the cycle by six hours, resulted in changes in the phase of activity in some foragers in the assay. The results shown in this study highlight the importance of temperature as a potential Zeitgeber in the field. Future studies will examine the possible functional and evolutionary role of the observed phase differences of circadian rhythms.

## Introduction

One of the major functions of endogenous circadian clocks is to maintain the appropriate timing (phasing) of physiological and behavioral processes with respect to daily variations (such as light-dark and temperature cycles) in the external environment. Typically, most of these clock-driven biological rhythms are not in perfect synchrony with such environmental cycles but, instead, occur at stable, fixed phases that anticipate or trail certain aspects (dawn, dusk, increasing or decreasing temperatures) of the environmental cycles(Moore and Rankin 1983, Frisch and Aschoff 1987, Panda et al. 2002, Hut and Beersma 2011). The endogenous circadian clock and the circadian rhythms orchestrated by it are thus entrained by daily environmental time cues (Zeitgebers). While circadian rhythms were originally measured as daily behavioral patterns, such as locomotor activity, egg laying, mating, and food acquisition, at the molecular level, circadian rhythms are driven by a set of proteins that generate negative feedback loops that regulate the transcription, translation and post translational modifications of a large number of canonical clock genes (Dunlap 1999, Blau 2001, Cyran et al. 2003, Gardner et al. 2006). These feedback loops generate near 24-hour (circadian) oscillations in the expression levels of the genes that make up the clock (Takahashi 1999, Ko and Takahashi 2006).

In honey bees, the circadian clock is associated with various complex behavioral processes that drive the survival and fitness of the colony. Drones and queens (the reproductive castes in the colony) mate at a specific time during the day (Galindo-Cardona et al. 2012). Foragers, who go out in search of different resources during daylight, learn the time and location of a food source and anticipate its availability on following days (von Frisch 1967, Moore 2001, Moore and Doherty 2009).Foragers also rely on the circadian clock for time-compensation, an essential component of their sun-compass navigation and dance language communication functions (Lindauer 1960, von Frisch 1967, Cheeseman et al. 2012) Although these and other processes rely on precise timing of the circadian clock, few studies have examined the potential environmental cues that entrain circadian rhythms in nature.

The first studies of honey bee circadian rhythms looked at foraging rhythms at the colony level (Buttel-Reepen 1900, Forel 1910, Wahl 1932, 1933). Studies examining the potential Zeitgebers that entrain circadian rhythms in honey bee colonies as well as individual bees concluded that light-dark cycles are the primary entraining agents (Renner 1960, Beier 1968, Beier and Lindauer 1970, Detrain et al. 1999). Studies in which groups of individuals from whole colonies were trained to visit a specific food source at a particular time of day were then translocated to a different time zone (ex. New York to California) exhibited foraging activity at the time relative to their original light-dark cycles. Over a span of several days, foragers from the transplanted colony re-entrained to the new light-dark regimen (Renner 1959). At the individual forager level, studies have demonstrated that light-dark cycles indeed entrain forager locomotor rhythms (Spangler 1972, Moore and Rankin 1985) and that the lights-off transition determines the forager’s phase of activity (Moore and Rankin 1993). However, the fact that honey bee colonies and individuals entrain to light-dark cycles does not exclude the possibility of entrainment by other environmental cues.

An important aspect of hive maintenance is that honey bees socially regulate colony temperature, keeping it optimally at 35°C (Heinrich 1980). At the behavioral level this is achieved by heat production via vibration of wing muscles, fanning (ventilation), and water evaporation inside the colony (Seeley 1974, Kronenberg and Heller 1982). Researchers examining the mechanisms underlying this thermoregulation found that individual variation at the genetic level is associated with differences in worker’s fanning response threshold to temperature (Jones et al. 2004). This variation results in different patrilines fanning at different temperatures and, therefore, promoting the stability of brood nest temperature (Jones et al. 2004). A primary challenge to colony thermoregulation is the daily variation in external heat associated with sunlight. The contribution of variation in circadian rhythms at the level of the individual bee to thermoregulation remains unknown.

Thermoregulation of honey bee colonies is essential for colony performance and survival (Heinrich, 1981; Heinrich, 1993). Experiments examining the effects of low temperatures (28-30°C) on brood development revealed that these temperatures can cause shriveled wings or other malformations, while brood kept at high temperatures (38°C-40°C) exhibit high mortality rates (Heinrich, 1993; Himmer, 1927). Subsequent studies showed that pupal development at 32°C, only 3°C lower than optimal core temperature, results in significant deficits in waggle dance behavior as well as learning and memory assays, compared to bees raised at 36°C (Tautz et al. 2003). Nonetheless, little is known about the effects of colony temperature on circadian rhythms in honey bees.

While in the tropics and neotropics environmental temperatures are somewhat stable throughout the year, honey bee colonies in temperate and subpolar regions are exposed to drastic temperature changes on an annual basis. Honey bee colonies are heterothermic. In the winter, temperature regulation is driven around the survival of the cluster. When a colony is exposed to a cold stress, its workers will cluster up densely in order to reduce colony heat loss(Southwick 1985). At the individual level, workers will produce heat by shivering their flight muscles in order to keep the core temperature of the cluster above the environmental temperature (Heinrich and Esch 1994, Stabentheiner et al. 2003). In contrast, during the spring and summer when colonies are rearing brood, temperature must be controlled within a very narrow range(Kronenberg and Heller 1982, Fahrenholz et al. 1989, Jones et al. 2004). Even small discrepancies from the 35°C optimal temperature for brood development can have negative consequences for adult worker fitness (Tautz et al. 2003, Jones et al. 2004).

Previous studies have explored the entrainment of circadian rhythms by square-wave temperature cycles and found that cycles with amplitudes greater than 9°C successfully entrain circadian locomotor rhythms of individual foragers in the laboratory (Moore and Rankin 1993). However, whether temperature cycles strong enough to entrain circadian rhythms exist inside the colony has not been thoroughly studied. Changes in environmental temperature in the laboratory have been shown to have strong effects on the endogenous period length of foragers (Fuchikawa and Shimizu 2007, Giannoni-Guzman et al. 2014). Furthermore, work from our group has revealed a broad range of individual difference in the endogenous period length of the circadian clock among foragers (Giannoni-Guzman et al. 2014). This large variation in the endogenous period length could be adaptive at the colony level with possible effects on fanning, shivering and clustering behaviors and could result in large differences in the phase of the circadian clock to time givers.

The primary goal of this study was to ascertain if temperature changes on a daily basis inside honey bee colonies and, if so, are these changes (temperature cycles) capable of influencing circadian rhythms of honey bee foragers? We measured light and temperature at various locations inside of the colony to determine if these potential Zeitgebers showed daily oscillations. We then explored the effects of temperature cycles observed in the colony on the circadian locomotor rhythms of honey bee foragers in the laboratory. Finally, we phase-shifted the temperature cycle in an attempt to confirm if the locomotor rhythms in individual bees were capable of stable entrainment to the temperature cycles.

## Materials and Methods

### Colony Light and Temperature Measurements

Light and temperature measurements were carried out using HOBO^®^ pendant data loggers (UA-002-64). Four loggers were placed inside the colony at the center, entrance, periphery and top, while one was placed outside the hive as shown (Figure 1A). The colony in which we collected these measurements was a two story colony in good health with a naturally mated queen laying eggs, and containing approximately 6-8 brood frames and 60,000 workers. With the exception of 30 minutes during July 11, 2013, when data were uploaded from the pendants and the batteries were replaced, temperature and light measurements were continuously collected in 30-minute intervals from June 13, 2012 until September 2, 2014. The bees and colonies described in our experiments were located in Gurabo, Puerto Rico. Throughout the year, mean high temperatures averaged from 28-30°C, while the low temperatures averaged between 16-20°C (Acevedo-Gonzalez et al. 2019, Feliciano-Cardona et al. 2020). Temperature and light data presented are averaged monthly values for the month of July 2013 (Figure 1). The peak to trough amplitudes were calculated using the calculated amplitudes of the temperature oscillations and multiplying by 2 (Figure 2). The phase of the average monthly temperature was calculated using cosine fitting in the circadian dynamics app (24 Dimensions LLC).

**Figure 1.**
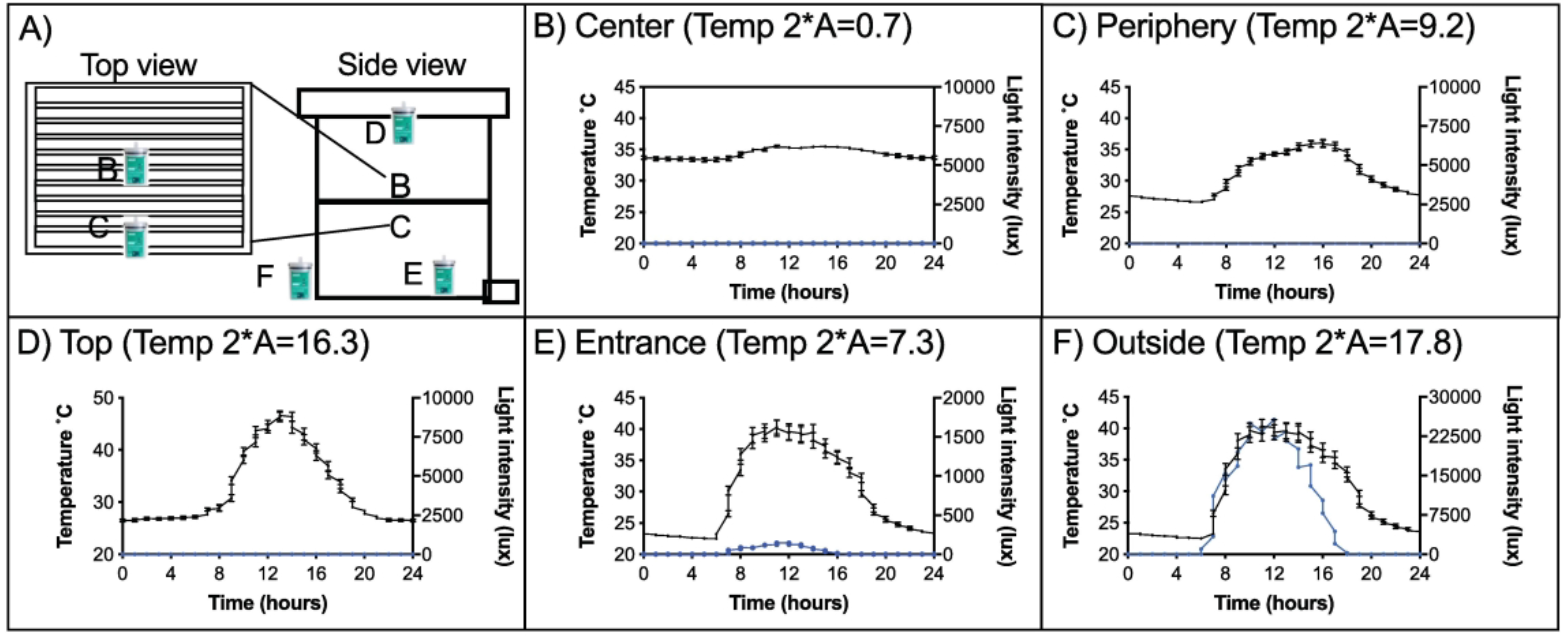
Temperature oscillates with a 24 hour period in the periphery of the colony. **A)** Top and side view of a two story honey bee colony presenting the positions of the 5 sensors used to measure environmental temperature and light, inside and outside the hive. Average temperature ±SEM and light environmental data logged at the **B)** center, **C)** periphery, **D)** top, **E)** entrance and **F)** outside the colony in the month of October 2013. Temperature was plotted on the left y-axis (black line), while light is plotted on the right Y-axis (blue line). As sensors move further away from the center of the colony peak to trough differences (Temp 2*A) of temperature oscillations increases from 0.7°C at the center of the colony up to 16.3°C in the top of the colony.

**Figure 2.**
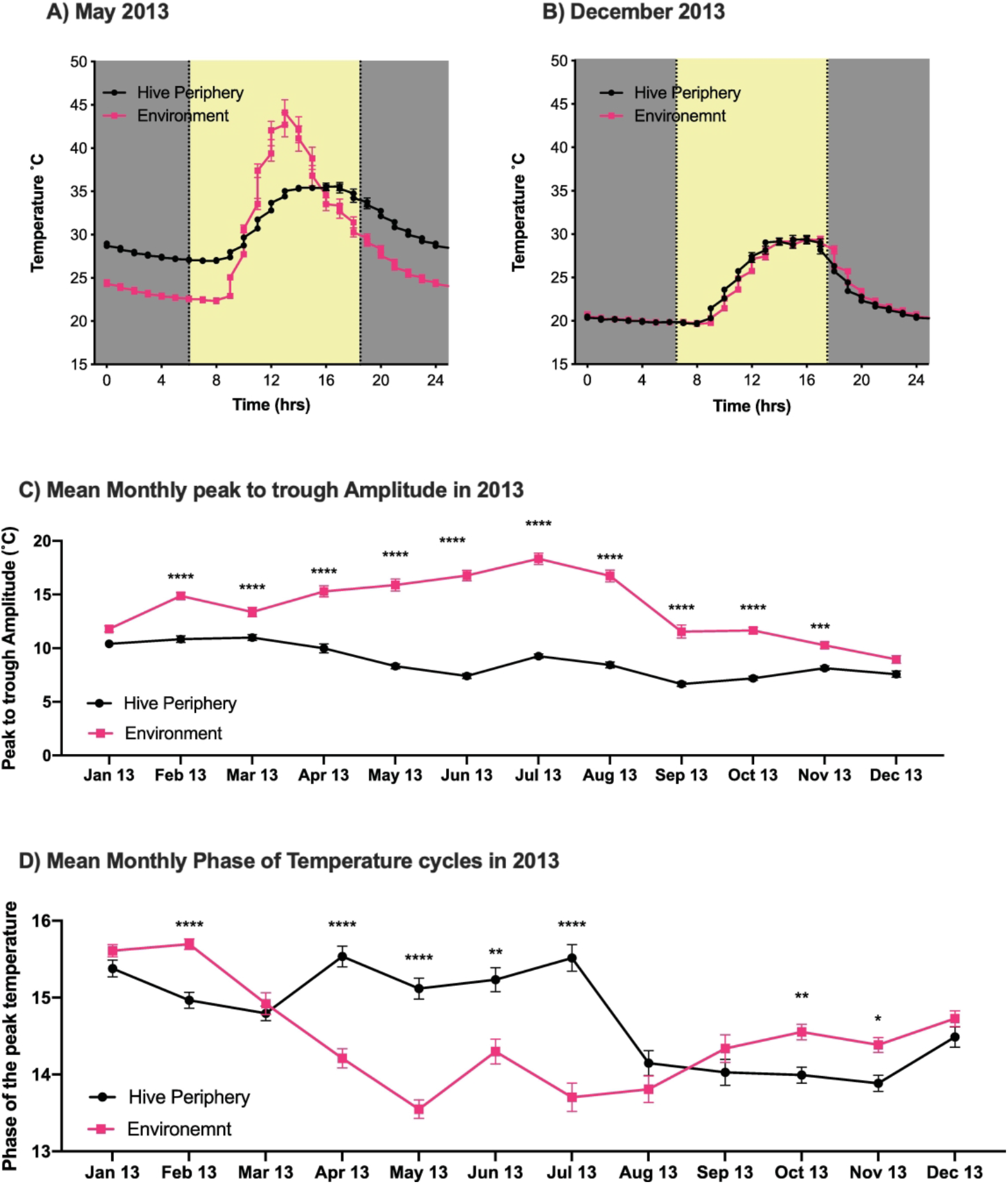
Bees actively regulate the phase and amplitude of temperature oscillations in the hive periphery. Average temperatures of hive periphery (black) and outside environment across the day in the months of **A)** May and **B)** December 2013. Grey and yellow in the background represent the photoperiods during each of these months. **C)** Mean ± SEM monthly peak to through Amplitude for the hive periphery and environmental monitors in 2013. Mixed effects model was significant time of year (F_(5.31, 316.2)_ = 47.11; *p<0*.*0001*\*\*\*\*) location of sensor (F_(11, 656)_ = 10087; *p<0*.*0001*\*\*\*\*) and their interaction (F_(11, 656)_ = 31.56; *p<0*.*0001*\*\*\*\*). Sidak’s multiple comparisons test between groups showed significant differences (*p<0*.*001\*\*\**) for each month with the exception of January and December. **D)** Mean ± SEM monthly phase of temperature cycles in the hive periphery (black) and the outside environment (pink). Mixed effects model was significant time of year (F_(7.369, 437.5)_ = 24.81; *p<0*.*0001*\*\*\*\*) location of sensor (F_(1, 653)_ = 25.07 *p<0*.*0001*\*\*\*\*) and their interaction (F_(11, 653)_ = 22.55; *p<0*.*0001*\*\*\*\*). Sidak’s multiple comparisons test between groups showed significant differences (p<0.01*) for marked months.

### Forager Collection

All of the bees utilized in our experiments came from colonies in good health that had naturally mated queens and were laying eggs at the time of the experiments. For each experiment, foragers were collected returning to the colony by blocking the entrance with 8-mesh wire screen and capturing them in tubes as previously described (Giray et al. 2007). Collected bees were provided with food and water during transportation to UPR Rio Piedras campus (30-40 min car ride). Once in the lab, bees were anesthetized and placed in locomotor activity monitors as previously described (Giannoni-Guzman et al. 2014).

### Experiment 1: Simulating temperature cycles of the colony in the laboratory

Locomotor activity recordings were performed inside an environmental chamber (Percival, I-30BLL), where temperature was programed to oscillate with an amplitude of 9.2°C, as observed in the periphery of the colony (Figure 1C). To ensure that the temperature was oscillating in the same manner (or as close as possible) as the observed oscillation in nature, the incubator was set up and calibrated 2 weeks prior to the experiment. Age was controlled by paint marking individual 1-day old workers and returning them to the colony on October 7^th^ 2014. Foragers were from the same colony of origin. Locomotor activity recordings began at 21 days of age and took place from October 28^th^ until November 7^th^ 2014. Phase analysis of locomotor activity was performed on days 6-11 of locomotor recording, to allow 5-6 days needed for bees to acclimate to square wave temperature cycles (Moore and Rankin 1993). Data of foragers in constant darkness at 35°C used is from previously published work (Giannoni-Guzman et al. 2014).

### Experiment 2: 6-hour phase advance of simulated temperature cycles

Foragers from the same colony were collected at the entrance of the colony September 18^th^, 2015 on a sunny afternoon and placed in constant darkness with oscillating temperature cycles later that same afternoon. Data shown and used for analysis were taken beginning the first midnight. Locomotor activity assays were conducted using the same environmental conditions as experiment 1 for the first 6 days of the experiment. On the 7^th^ day the environmental temperature cycle was advanced by 6 hours and was kept with this timing until the end of the experiment (Figure 4 and 5).

**Figure 3.**
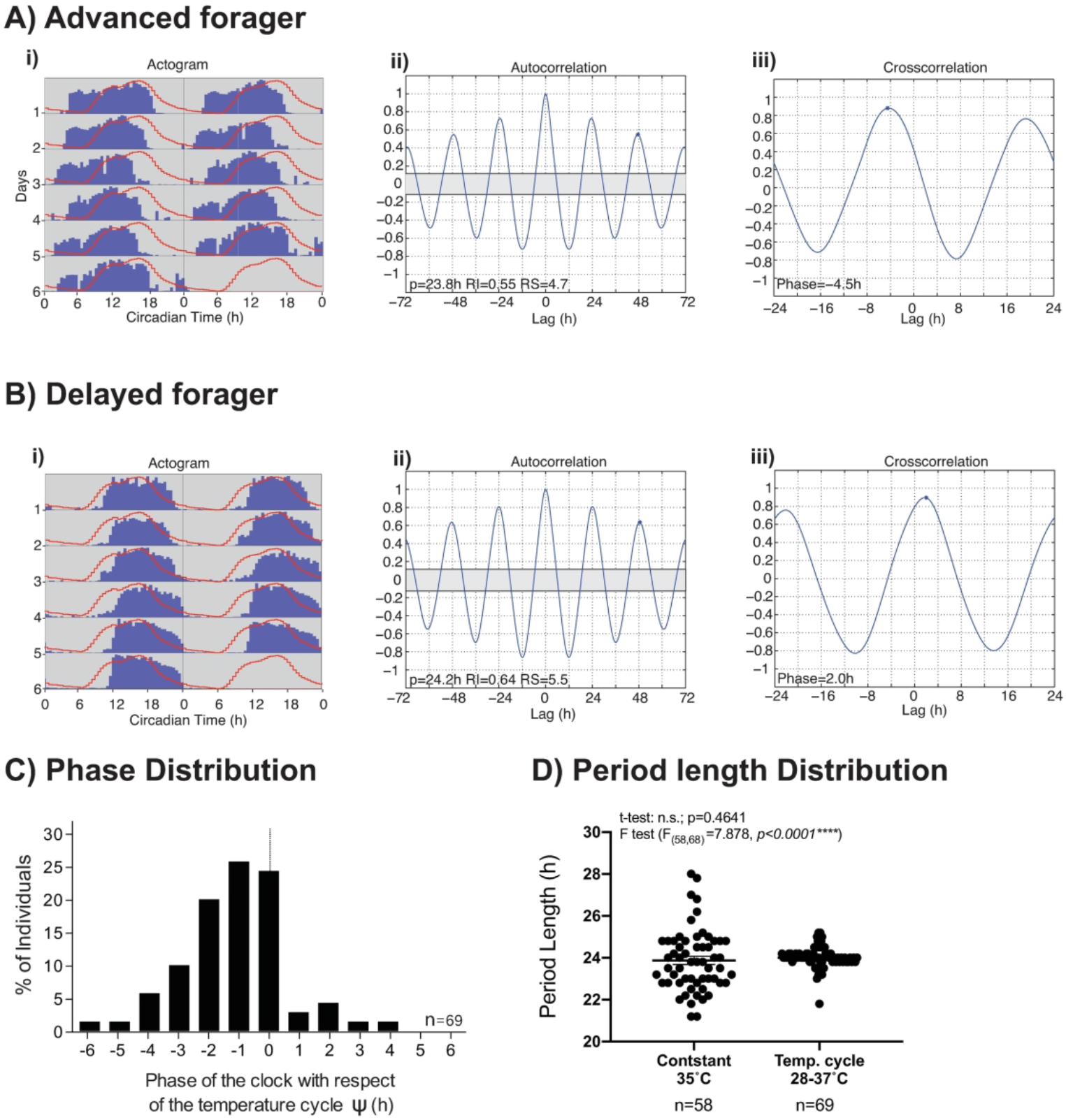
Simulating peripheral temperature oscillations is associated with strong rhythmicity and large individual differences in circadian phase. Representative examples of foragers that presented **A)** phase advance or **B)** phase delay with respect to the simulated temperature cycles in controlled laboratory conditions. **i)** Double plotted actogram of representative individual locomotor activity (blue bars) with simulated temperature overlayed (red lines). **ii)** Autocorrelation analysis was utilized to determine period length and overall rhythm strength of locomotor rhythms (Levine et al., 2002). **iii)** Phase of the locomotor rhythms with respect to the temperature cycles (*ψ*) was quantified using cross-correlation analysis, which compares the distribution of locomotor activity with the temperature cycle and yields the phase as the lag (h). **C)** Frequency distribution of the phase of locomotor activity with respect to temperature cycle. **D)** Period length distributions of foragers kept under constant conditions (darkness, 35°C) and bees exposed to the measured peripheral temperature oscillations (t-test: n.s.; p=0.4641) (F_(57,68)_=7.878, *p<0*.*0001\*\*\**).

**Figure 4.**
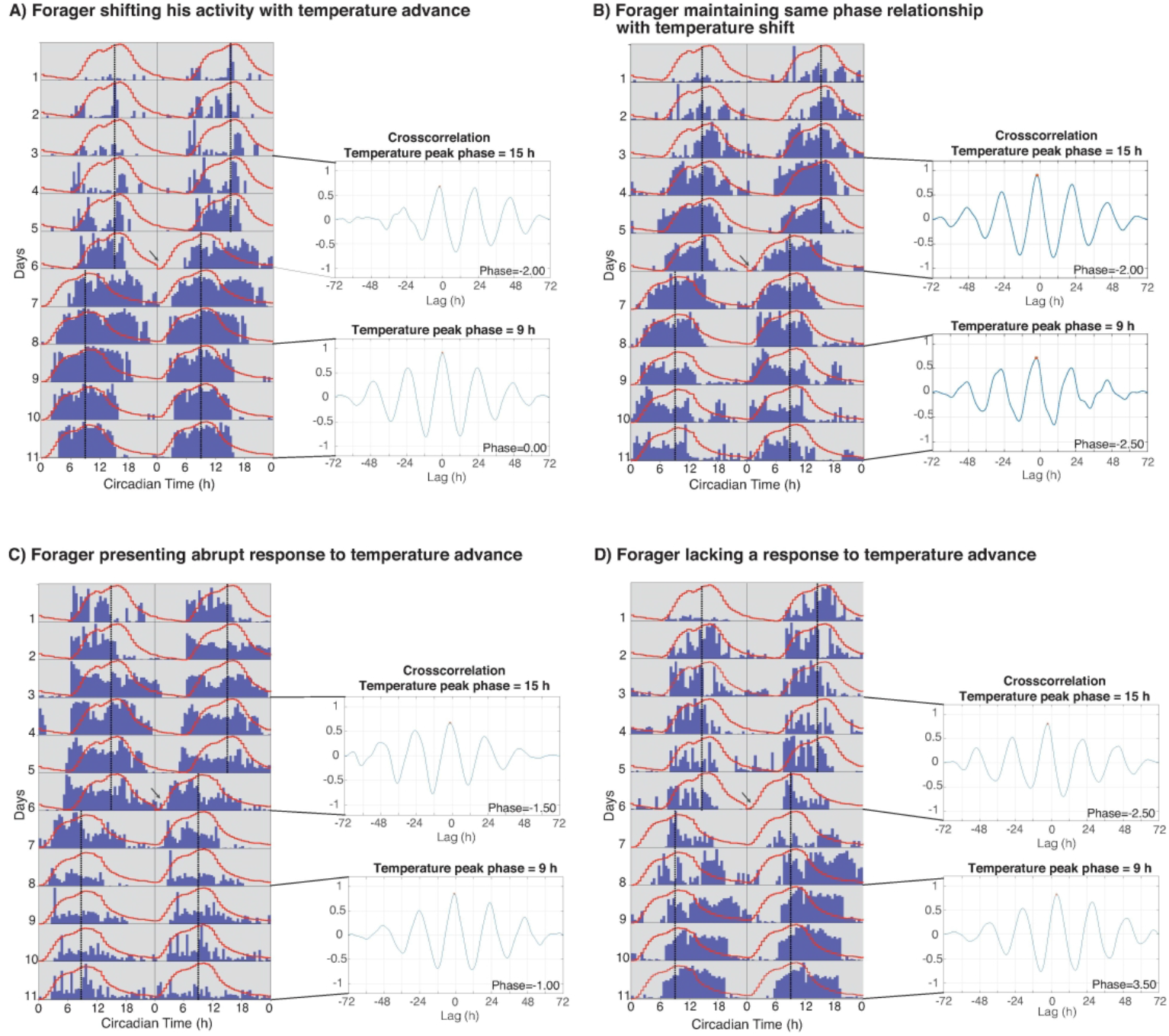
Simulated temperature oscillations are capable of phase advancing the locomotor activity of forager bees. Representative double plotted actograms of locomotor activity (blue bars) with simulated temperature cycles overlayed (red lines). At midnight on the 7^th^ day the temperature cycle was advanced 6 hours. For each activity plot the phase of locomotor tor rhythms with respect to the temperature cycles (*ψ*) was quantified using cross-correlation analysis. Two cross correlations were calculated, the first on days 4, 5 and 6 before the temperature manipulation and the second 48 hours after temperature change (days 9, 10 and 11). **A)** Representative plot of forager synchronizing his locomotor activity rhythm to the phase of temperature in response to temperature advance. **B)** Representative individual maintaining a consistent phase relationship with simulated temperature cycles before and after the temperature manipulation. **C)** Representative plot of forager showing an abrupt response to temperature advance. **D)** Representative forager showing no response to temperature advance.

**Figure 5.**
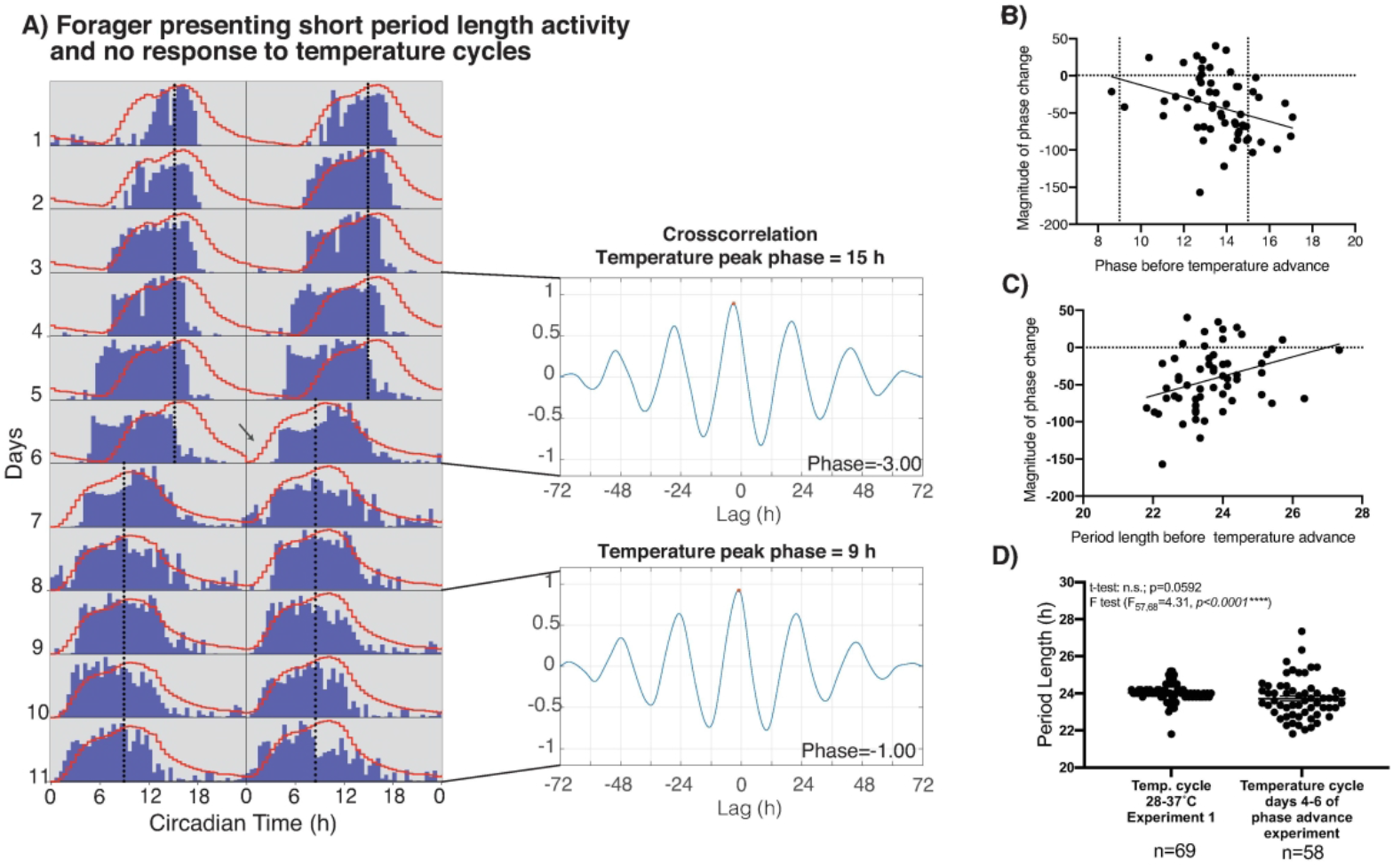
Relationship of phase changes and period length of locomotor activity. **A)** Representative double plotted actogram of forager presenting short activity period length and no apparent response to temperature cycle manipulation. Cross correlation analysis for days 4, 5 and 6 before the temperature manipulation and days 9, 10 and 11 are shown with respect to the temperature cycle during these days. **B)** Correlation examining the relationship between the magnitude of phase change and the phase before the temperature advance resulted in a significant correlation (Pearson *r*(58)=−0.3282, p=0.012*). Magnitude of the phase change was calculated as follows: ((Phase after temperature advance - Phase before temperature advance)/6) ×100. **C)** Correlation of the Magnitude of the phase change and the period length of the locomotor activity before the temperature advance (Pearson *r*(58)=−0.3411, p=0.009**) **D)** Period length distributions of foragers kept exposed to temperature cycles from our first experiment and days 4-6 before the environmental manipulation of phase advance experiment (t-test: n.s.; p=0.0592) (F_(57,68)_=4.31, *p<0*.*0001\*\*\*\**).

### Data analyses

Locomotor activity data were processed using MatLab® toolboxes developed in Jeffrey Hall’s laboratory (Levine et al. 2002). Period length was calculated using autocorrelation analysis. The phase angle (ψ) of the acrophase locomotor rhythm in relation to the acrophase temperature cycles to which bees were exposed was determined via cross correlation analysis. Changes in the phase of temperature measurements throughout the year were analyzed via Mixed-effect Modeling and post hoc tests were performed between groups.

## Results

Previous work has shown that square-wave temperature cycles are successful Zeitgebers for the circadian clock of honey bees under laboratory conditions (Moore and Rankin 1993). However, whether temperature cycles capable of entrainment of the circadian rhythm of honey bees are present inside the colony was unknown. To examine if temperature cycles are present in honey bee colonies, we placed 5 data loggers at different locations of the colony (center, entrance, top, periphery and outside) and recorded light and temperature measurements in 30-minute intervals (Figure 1A). Our results revealed that temperature oscillates inside honey bee colonies in a daily manner, with amplitudes greater than 7 degrees (Figure 1). Specifically, we observed that at the center of the colony temperature was maintained within 35±0.7 °C, while daily temperature cycles at the entrance (peak to trough amplitude=7.3°C), periphery (peak to trough amplitude=9.2°C) and top (amplitude= 16.3°C) of the colony were detected. The control data logger that was placed outside the colony, as expected, detected strong temperature and light-dark cycles with greater amplitude than those detected in the colony (Figure 1E). In addition, the presence of light inside the colony was only detected by the logger at the entrance and was 200 times less at its highest peak than that detected by the logger outside the hive (Figure 1E, F).

Having observed that temperature oscillates in parts of the colony, we next looked at whether there were significant seasonal changes in these temperature oscillations inside the colony. We focused on comparing the peripheral temperature (Figure 1C) with that of the outside environment. Our results show that the amplitude and the phase of temperature cycles at the periphery of the hive varies throughout the year. We observed significant phase and amplitude differences between the periphery and environment during the spring and summer months, for example May (Figure 2A), with similar phasing and amplitude of the cycles in the month of December (Figure 2B). The amplitude of the temperature cycles in the periphery of the colony was significantly different from the environment for all months with the exception of December and January (Figure 2C). Importantly, the variance over the year was significantly less inside the periphery of the hive than outside (F_(11,11)_=3.89, *p=0*.*0334\**). Comparing the mean phase of the two temperatures revealed significant phase delays in the peak of peripheral temperature cycles from April to July compared to those of the environment (Figure 2D). These data suggests that bees inside of the colony actively regulate the phase and amplitude of temperature oscillations in the periphery of the hive throughout the year.

Given the previously reported effects of constant environmental temperature on the endogenous rhythm of honey bees (Fuchikawa and Shimizu 2007, Giannoni-Guzman et al. 2014), we examined the possible effect of the observed temperature cycles on the locomotor rhythms of foragers. We simulated the temperature cycle recorded from the periphery of the colony in the laboratory and measured the locomotor activity rhythms of individual foragers subjected to this cycle under constant dark conditions. We simulated the peripheral temperature oscillation of the month in which we performed the bees were collected (October), which had a 9.2° C peak to trough amplitude. We hypothesized that exposing foragers to simulated temperature cycles would result in their locomotor activity rhythms achieving a stable, consistent phase relationship with the temperature cycle. Alternatively, locomotor activity could still be influenced by the temperature cycle but not attain a fixed phase relationship. Our results showed consistent phase relationships, but with a broad range of individual differences among the phases (*ψ*) of locomotor activity with respect to the temperature cycle (Figure 3).

Most individuals in our sample were phase advanced (Figure 3A), while some showed a delayed phase (Figure 3B). Phase quantification for each individual was performed via cross-correlation of the locomotor activity and the temperature cycle (Figure 3, panel (iii)). Frequency distribution of the phase of individuals revealed that more than 60 percent of foragers are phase advanced to the temperature cycle (Figure 3C). In addition, the mean period ± SE of the activity rhythm under the temperature cycle was 24.00 ± 0.06 h and its variance was significantly smaller than that of the period of foragers under constant conditions (Giannoni-Guzman et al. 2014) (Fig. 3D). The 24.00 hour period of the locomotor activity rhythm under the temperature cycle, coupled with the fact that the variance in period is smaller under the temperature cycle compared to constant conditions (23.8 ± 0.19 h) (Giannoni-Guzman et al. 2014), suggests that the locomotor activity rhythm is entrained to the temperature cycle.

To further test if the observed temperature cycles were capable of entraining the locomotor rhythms of foragers, we performed a 6-hour phase advance of the temperature cycle on the 7^th^ day of locomotor activity measurements (Figure 4). We hypothesized that if the temperature cycles observed in the colony were capable of entraining the circadian locomotor rhythms, then shifting the temperature cycle would result in a shift in the phase of locomotor activity of bees over several days (transients) until resuming the previous phase relationship with respect to the temperature cycle. Consistent with this hypothesis, we did observe that approximately 51% individuals showed a response to the phase advance of the temperature cycles (Figure 4A, B and C). Within these individuals 56% gradually advanced their locomotor phase of activity (Figure 4A-B), consistent with the expression of transients associated with re-entrainment, while 44% abruptly advanced their phase with respect to the temperature cycle, which could be consistent with masking (Figure 4C). In these cases, the activity apparently moved to establish a consistent phase relationship with the new temperature cycle.

In addition to the foragers that responded to shifts in the temperature cycle, we also observed that approximately 49% of foragers were apparently unaffected by the temperature shifts (Figure 4D). Furthermore, a large subset of the foragers in this experiment exhibited short-period activity rhythms and no discernable response to the temperature advance (Figure 5A). To determine if the magnitude of the shift correlated with the phase prior to the temperature advance, we calculated the magnitude of phase changes using the following formula: ((Phase after temperature advance - Phase before temperature advance)/6) x100. This correlation revealed that the later the peak of activity prior to the shift the greater the phase change (Figure 5B). However, a correlation of the magnitude of phase change and period length before the shift was positive (Figure 5C), suggesting changes in phase are mostly driven by changes in the period length of the foragers examined. Comparing the period length distribution of foragers from our previous temperature cycle experiment with that from the days prior to the phase advance of this experiment revealed significant differences in variance between the experiments (Figure 5D). The fact that the phase advance experiment shows greater variance in period lengths, suggests that many bees were free running and therefore unresponsive to the temperature (Figure 4D and Figure 5A). This period length distribution is similar to the previously observed difference in variance when comparing foragers under constant conditions to those exposed to a temperature cycle (Figure 3D). However, although variance was greater it was still significantly less than that under constant conditions (F_57,57_=1.83, *p=0*.*0250\**). Taken together, our results suggest that while 51% bees responded to temperature and shifted their activity in the direction of the temperature cycle phase shift, 49% of bees in this experiment did not respond to either of the temperature cycles.

## Discussion

Here we show that at the periphery of the colony, where foragers spend much of their time (Van Nest et al. 2016), temperature significantly oscillates in a 24 hour period (Figure 1). The amplitude and phase of these oscillations change with respect to the time of the year, and the amplitude varies even less than the environmental temperatures (Figure 2). These findings indicate a tight regulation of temperature oscillations in the periphery of the hive. Simulating this temperature signal in the laboratory is sufficient to synchronize and phase advance the circadian locomotor rhythms of some forager bees, suggesting that temperature could be an important Zeitgeber in the colony (Figure 3, 4, 5). Interestingly, we found that there are large individual differences in the phase of locomotor rhythms with respect to the temperature cycles as well as the responses to a temperature phase shifts. Taken together, we believe that temperature is an important environmental cue inside the colony capable of entraining the circadian rhythms of foragers in nature.

Until recently, the circadian clock of honey bees was thought to be entrained mainly by light-dark cycles (Renner 1959, 1960, Moore and Rankin 1985). Recent work has shown that social cues, such as substrate-born vibrations and colony volatiles are capable of entraining and synchronizing the circadian clock of bees (Bloch et al. 2013, Fuchikawa et al. 2016, Siehler and Bloch 2020). However, these experiments do not take into account temperature changes that occur within the colony. Our findings in the present study, as well as those from previous studies (Kronenberg and Heller 1982), suggest that temperature cycles, strong enough to synchronize the circadian rhythms of foragers, are present inside the colony (Figure 1).

The seasonal changes we observed in peripheral temperature (Figure 2), suggest that the active regulation of temperature inside the colony is not limited to the brood nest. This maintenance of the oscillation could serve as a possible entrainment cue for times of the year when bees are unable to go out, such as winter. The delayed phase of the temperature peak during the spring and summer months could potentially play an important role in circadian synchronization for bees inside the colony during their more active periods of the year. Future studies will examine the role of these temperature oscillation in swarming, timing of drone flights and overwintering preparations inside the colony.

Previous research under laboratory conditions proposed that light and temperature may have a synergistic effect on the circadian clock system in honey bees (Moore and Rankin 1993). Experiments employing both light and temperature cycles show that some individuals responded most strongly to the presence of light, while others concentrated their present locomotor activity during times of overlap between the photophase and higher temperatures (Moore and Rankin 1993). Since foragers have been shown to stay inside the colony after visiting their food source (von Buttel-Reepen 1903, Körner 1940, von Frisch 1940, Moore et al. 1989, Seeley 1995), it is possible that, while inside the dark colony, foragers rely on temperature cycles to accurately maintain entrainment to the natural day-night cycle and use light input as a Zeitgeber when foraging. This hypothesis stems from the lack of light inside most of the colony (Figure 1). In order to test this hypothesis, further studies exploring the functional role of temperature cycles in the colony as well as the mechanisms of circadian entrainment to temperature cycles are needed.

Consistent with the results from the Moore and Rankin (1993) study, we observed that honey bee foragers synchronize to these simulated peripheral temperature cycles. To our knowledge, this is the first time that gradual temperature cycles, simulating those observed in the field, have been utilized in the laboratory. In addition, by simulating the peripheral temperature cycles we found that foragers present a large degree of individual variation in the phase of locomotor rhythms with respect to temperature (Figure 3). Although further studies are needed to understand the functional and evolutionary role of this variation, we can speculate that phase differences of the individual honey bee foragers may result in differences in the temporal allocation of tasks. The individual differences within the foraging population could potentially help with smoothing temperature regulation processes of the colony. This idea is consistent with work where decreasing the genetic diversity of the colony decreases the colonies ability to regulate temperature (Jones et al. 2004, 2007). Another possible functional role for this variation is foraging specialization. This hypothesis would be in line with the results of a recent study that suggests the existence of shift work in honey bees and its genetic component (Bernhard Kraus et al. 2011, Giannoni-Guzmán 2016, Giannoni-Guzmán et al. 2020). Alternatively, the observed differences in the phase of circadian rhythms could be the result of previous entrainment to an outside stimulus, such as light, nectar or pollen. The latter can be evidenced by the ability of foragers to successfully be trained for several days to a food source (Frisch and Aschoff 1987).

When performing a 6-hour advance on the temperature cycle, we observed that approximately 51% individual’s locomotor activity advanced and reached stable synchronization with the new temperature cycles, while 49% were unaffected by either temperature cycle (Figure 4). Within individuals unaffected by temperature, we observed a large number of individuals with short period length. Surprisingly, most of the foragers in this experiment presented short locomotor activity period lengths (Figure 5). This result differs from our previous experiment (Figure 5D) and could be the result of the time of year the experiments took place and that a different colony was used for each experiment. This variation in the change of locomotor activity patterns may reflect differences in the perception of temperature by the circadian system of foragers. Future work could simulate temperature cycles of the higher amplitudes cycles found inside the hive and test bees response to temperature in greater detail

Our results show evidence for temperature entrainment in the form of a strong response to phase advance of temperature by some foragers (Figure 4). Furthermore, given the large degree of individual variation in the free-running period of honey bees and their tendency to present short periods, we interpret the proximity to 24 hour periodicity and the significant decrease in variance as an additional sign of entrainment (Figure 3). Although some individuals gradually shifted their activity after the phase advance, as expected of circadian entrainment, there were individuals that shifted abruptly (Figure 4C). Further experiments to determine whether this particular groups response to temperature represents entrainment or masking are required. Specifically, exposing foragers to temperature cycles and later transferring them into constant conditions would clarify if abrupt responses to phase advance are masking.

Although the mechanisms for light and temperature input to the clock of honey bees remain to be elucidated, studies in *Drosophila* have shown that temperature cycles successfully entrain the locomotor and molecular rhythms (Tomioka et al. 1998, Yoshii et al. 2002, Glaser and Stanewsky 2005, Boothroyd et al. 2007). Light and temperature act in a synergistic manner to entrain both locomotor activity and molecular clock of *Drosophila* (Yoshii et al. 2009). Furthermore, studies indicate that there are clock cells in the brain that selectively respond to temperature cycles, while other clock cells respond to light/dark cycles (as reviewed by Ki et al. 2015). Given the similarities between the neural clock of bees and *Drosophila* (Fuchikawa et al. 2017, Beer et al. 2018), it is likely that at least some of the properties uncovered in the fly with respect to the clocks response to temperature will be analogous in the bee circadian clock.

While our results show how temperature variation in honey bee colonies can synchronize the circadian rhythms of foragers, other environmental factors remain to be considered. For instance, we have observed that humidity in the periphery of the colony also oscillates in a 24 hour cycle and changes seasonally (unpublished results). In plants it is clear that humidity is capable of entraining the circadian clock (Mwimba et al. 2018). However, whether these oscillations are capable of entraining the clock of bees remains a subject of future research.

Taken together, our results indicate that temperature cycles in the colony are capable of synchronizing locomotor rhythms of honey bee foragers. Individual differences in the response to phase advances suggest differences in the sensitivity of the clock to changes in temperature. Future studies will explore the importance of temperature as a time giver in typical colony conditions and its synergistic effects with light and other social cues. Individual differences in the phase of circadian rhythms are loosely reminiscent to those of circadian chronotypes in human populations and with further studies, honey bees could be a potential model for these differences in human populations. Our study adds to the rich and complex interactions of temperature and social organization in honey bees, demonstrating altered temperature effects on circadian rhythms. Research into the functional relevance of this synchronization and the seasonal differences in the colony temperatures may lead to a better understanding of the evolutionary relation of circadian rhythms and sociality.

## Acknowledgements

We would like to thank Dr. Arian Avalos, for help with the experiments. Thanks Dr. Luis de Jesus for their comments and suggestions. We would also like to recognize the director, Manuel Diaz and the personnel of the Gurabo Experimental Agriculture Station of the University of Puerto Rico at Mayaguez for use of facilities at ‘‘Casa Amarilla’’. We would like to thank Josue Rodriguez for his help with data processing. This work was sponsored by the National Science Foundation (NSF) awards 1026560, 1633184, 1707355 and the National Institute of Health (NIH) 2R25GM061151-13, P20GM103475.

